# Role of educational attainment, cognitive performance and intelligence in neurodegeneration: a bidirectional Mendelian randomization study

**DOI:** 10.1101/692855

**Authors:** Sandeep Grover, International Age-related Macular Degeneration Consortium (IAMDGC)

## Abstract

I examined the potential bi-directional causality between educational attainment (EA) (n = 766,345) and age related macular degeneration (AMD) (cases (n) =16144, controls (n) =17832) using the summary GWAS datasets on individuals with European ancestry. I used datasets on other late-onset neurodegenerative diseases including Alzheimer’s disease (AD) and Parkinson’s disease (PD) as controls to validate the findings. A risky effect of EA on AMD was observed (OR=1.318, 95% CI=1.080 −1.610, P=0.0068) after ruling out potential pleiotropy and absence of reverse causality. I further replicated previously observed protective and risky causal associations of EA with AD and PD.

## Introduction

Age related macular degeneration (AMD) is a late onset neurodegenerative disease that affects the retina region of the eye^1, 2^. It is also the third most leading cause of permanent vision loss. The etiology of AMD is poorly understood with several risk factors identified through large scale observational studies^3^. In addition to age and family history of AMD, cigarette smoking, obesity and hypertension are believed to be the most commonly observed risk factors for predisposition to AMD^4^. Furthermore AMD has also been shown to be associated with increased risk for cognitive impairment including Alzheimer’ disease (AD)^5^.

Recently potential role of educational attainment in neurodegeneration has aroused considerable interest possibly by modulating the cognitive performance (CP) ^6, 7^. The enhanced CP is believed to provide protection against AD and may reduce motor deficits in Parkinson’disease (PD). Furthermore, the modulatory role of educational attainment on neurodegeneration was recently confirmed by Mendelian randomization (MR) studies employing large scale GWAS studies to elucidate potential causality^8, 9^.

The MR employs genetic variants as randomized proxy markers of risk factors, also known as genetic instruments to check for directionality in a relationship between a risk factor such as educational attainment and a disease or outcome like AD or PD^10^. Using the largest cohorts on AD and PD available to date, educational attainment was most recently shown to play a protective casual role in AD and risky casual role in PD^8, 9^. Henceforth, this motivated me to explore the potential causal role of educational attainment in AMD.

## Materials and Methods

I conducted a two-sample MR study employing summary level estimates to explore the causal role of EA on AMD. I identified genetic instruments that influence EA using complete summary GWAS datasets provided by Social Science Genetic Association Consortium (SSGAC) from their recent publication^11^. I performed clumping on SNPs associated with EA (p-value< 5×10^−8^) using the Two Sample MR package (version 0.4.22) in R (version 3.4.4) to identify completely independent loci (r^2^ = 0.001) (http://cran.r-project.org/).

The summary estimates of the identified genetic instruments for the outcome dataset on AMD were extracted from the discovery cohort of a recent meta-analysis of GWAS on 16,144 AMD cases and 17, 832 controls of European ancestry^12^.

If SNPs were not available in the AMD GWAS dataset, proxy variants were identified using an R^2^ cut-off of 0.9. I used F-statistic to judge the strength of the genetic instruments, and power calculation was done using the online tool available at http://cnsgenomics.com/shiny/mRnd/). I used inverse variance weighted (IVW) fixed effect method as primary method to do causal estimate analysis and heterogeneity analysis including use of additional MR methods was conducted as described elsewhere^13^.

A Bonferroni corrected threshold of P=0.0167 was considered to be significant, considering the primary association analysis of EA with PD, AD and AMD. The selection of appropriate genetic instruments and validity of analysis was further verified by replicating the previously reported causal association of EA with AD and PD^8, 9^.

As a part of sensitivity analysis to further check the strength of my results, the causal analysis was repeated by excluding variants with potential confounding variables (cigarette smoking, obesity, hypertension, INT, and CP) identified through the Phenoscanner database (http://www.phenoscanner.medschl.cam.ac.uk/). As a part of secondary analysis, I further conducted MR analysis using genetic instruments for highly correlated phenotypes such as CP and INT to rule out role of EA as a potential confounder in association analysis with AMD^11, 14^. And lastly, I conducted check for potential causal role of genetic predisposition to AMD, AD and PD on educational EA, INT and CP to confirm the directionality of earlier observed direct causal associations.

## Results

Descriptive statistics of the GWAS datasets and prioritized genetic instruments used in the present manuscript are listed in **Table 1**. Furthermore, the summary data used for all the analyses (primary, sensitivity, and secondary) in the current article has been further provided in the **Supplementary Table 1**. A total of 318 genetic instruments were prioritized used the EA GWAS dataset on 766, 346 individuals. The SNP id rs77719387 or its proxy was not available in the AMD dataset leading to finalization of 317 SNPs available for the causal analysis with all of them fulfilling the F-statistics criteria >10. The casual association analysis of EA with AMD has been further provided in **Table 2**.

**Tablel 1.**
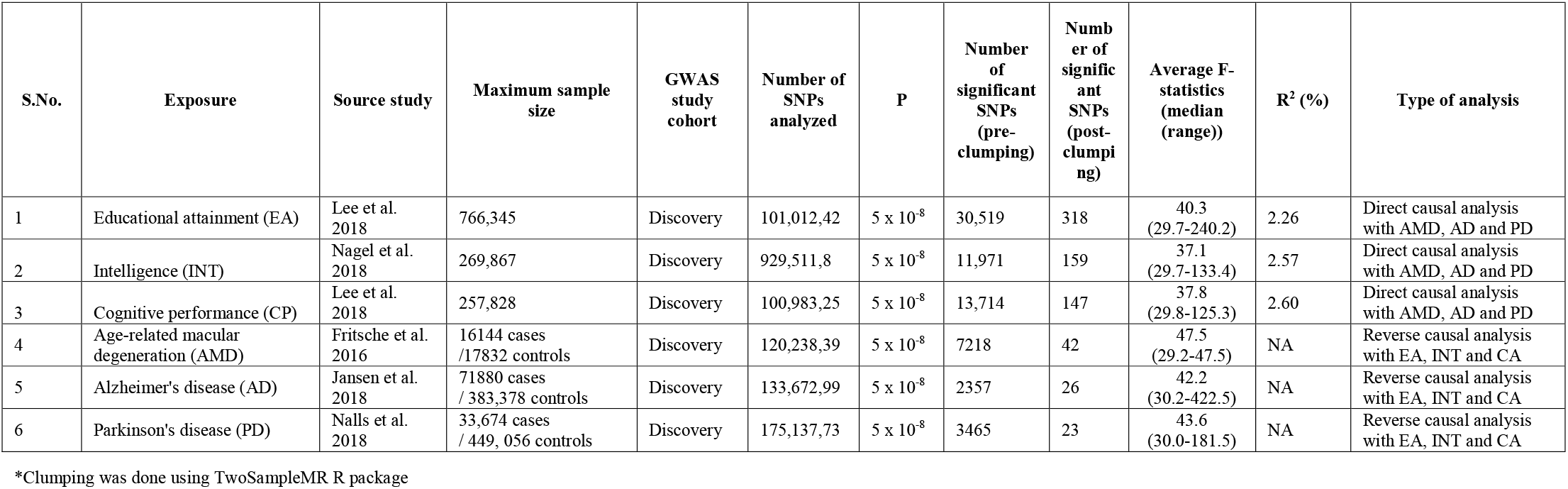
Details of discovery GWAS datasets explored and prioritized instruments used for the main analysis in the present study.

**Tablel 2.**
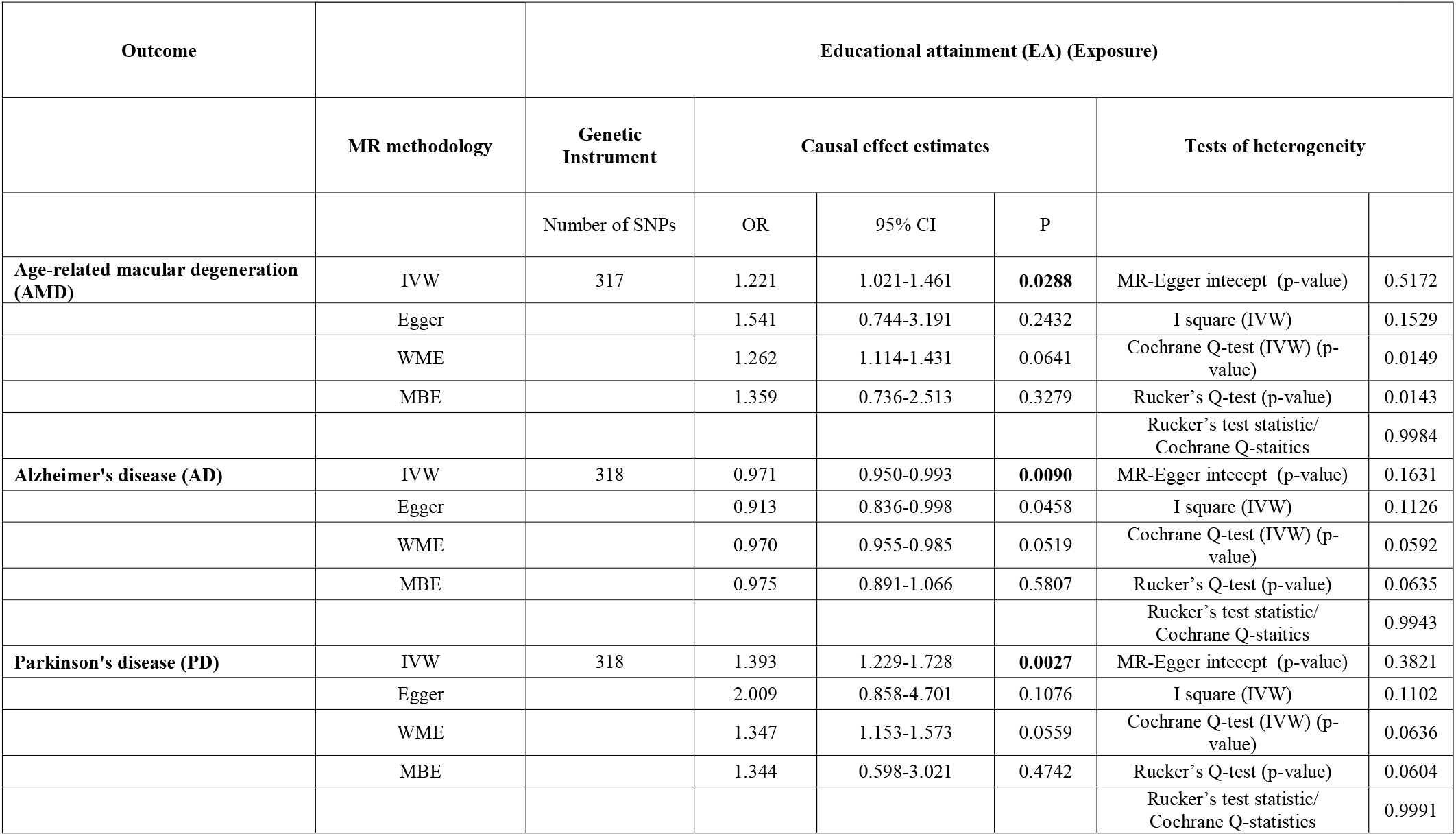
Causal effect estimates using different Mendelian randomization methods and heterogeneity analysis of causal effect estimates for Age-related macular degeneration (AMD), Alzheimer’s disease (AD), and Parkinson’s disease (PD) using educational attainment (EA) as exposure.

I observed a risky causal effect of EA on AMD (OR=1.221, 95% CI=1.021-1.461, P=0.0288) in the absence of significant pleiotropy (MR-Egger P=0.5172). A similar trend was observed using Weighted median method (OR=1.262, 95% CI=1.114-1.431) which accounts for undetected pleiotropy. I further identified a total of 57 genetic instruments as potential pleiotropic variants based on previously reported association of loci represented by these genetic instruments with known risk factors of AMD (**Supplementary Table 2**). The exclusion of these SNPs further strengthened my results (OR=1.318, 95%CI=1.080-1.610, P=0.0068). I further used the same genetic instruments to replicate previously reported causal association with AD (cases (n) = 71,880; controls (n) = 383,378) and PD (cases(n)=33,674; controls(n)=449, 056). A protective association of EA with AD (OR=0.971, 95% CI=0.949-0.992, P=0.0090) and a risky association of EA with PD (OR=1.393, 95%CI=1.122-1.728, P=0.0027) were observed. A total of 38 genetic instruments were available as proxy markers of AMD for checking causal role of AMD on EA. I observed absence of effect of AMD on EA, thereby further strengthening the observed causal association on EA on AMD (OR=1.003, 95%CI=0.998-1.009, P=0.2197, MR-Egger P=0.8446). Similarly, I observed lack of association of proxy markers of AD and PD with EA (Data not shown here), thereby suggesting that genetic predisposition to neurodegeneration has not role on EA.

Lastly, the bidirectional MR analysis was repeated using genetic markers of phenotypes closely correlated to EA namely INT (n=269,867) and CP datasets (n=257, 828) on AMD, AD and PD (**Supplementary Table 3**). I did not observe any association of INT and CP with AMD in both direct and reverse causal analysis. However, a moderate protective and risky effect of CP was observed in predisposition to AD and PD suggesting presence of residual confounding in the association of EA with AD and PD.

## Discussion

This is the first comprehensive study studying influence of EA, INT and CP on late-onset neurodegeneration using a bidirectional MR study design. To the best of my knowledge, this is the first MR study reporting causal role of EA on AMD. The casual association was further retained after excluding genetic variants known to be associated with potential risk factors of AMD.

Few observational studies have earlier explored the association between EA and correlated phenotypes on AMD. The Beaver Dam Eye study was amongst the earliest to show a relationship between education and incidence of AMD in an American population comprising of 3681 adults. The study showed that individual with 16 or more years of education at a higher risk of developing AMD compared to those with fewer than 12 years of education^15^. A population based cross sectional study of 3280 Asian individuals (Singopore Malay population) comprising of 168 early AMD had earlier demonstrated that individuals with low educational level were more likely to have early AMD (OR = 2.2; 95% CI = 1.2 – 4.0)^16^. Another recent study in Asian population (Chinese) comprising of 357 AMD patients showed increase prevalence of AMD among individuals with lower education level^17^. It is quite evident that most of these studies were underpowered and potential undetected confounding cannot be ruled out.

On the contrary, my MR study employed approx. 800,000 individuals including 16,144 individual with AMD to draw the conclusion in the absence of confounding. My study however had few limitations. Firstly, I could not use complete GWAS datasets on EA and PD, due to nonavailability of 23andMe datasets. This forced me to prioritize genetic instruments using a subcohort of datasets used in the published manuscripts. This approach helped me to conduct a bidirectional MR using the same datasets and hence avoiding potential bias in making interpretation on causal association. Nevertheless, loss of small proportion of individuals is unlikely to have any effect on the reported associations. Furthermore, large number of genetic instruments with high average F-statistics and replication of previously observed causal association’s demonstrated validity and strength of my results. Furthermore, the findings need to be replicated in the Asian population which is believed to be at decreased risk to AMD compared to European population.

The consistent role of EA on influencing the neurodegeneration in the absence of role of other closely related proxy markers including INT and CA suggests the need for conducting a systematic MR using both correlated exposures and outcomes with potential pathophysiological overlapping mechanism^18^.

---

**Supplementary table 1.** Harmonized summary effect estimates from exposure and outcome datasets used for the conduct of Mendelian randomization (MR).

**Supplementary table 2.** List of variants identified as pleiotropic variants in the association analysis of educational attainment (EA) with Age-related macular degeneration (AMD).

**Supplementary table 3.** A summary of direct causal association analysis of educational Intelligence (INT), and cognitive perfrormance (CP) with Age-related macular degeneration (AMD), Alzheimer’s disease (AD) and Parkinson’s disease (PD).

## Acknowledgement

I thank Prof. Inke König for providing the institutional facilities and research environment for conduct of the research. I acknowledge the investigators of the Social Science Genetic Association Consortium (SSGAC) for sharing the summary statistics from GWAS on EA and CP, Center for Neurogenomics and Cognitive Research-Complex Trait Genetics Lab (CNCR CTGLAB) for sharing the summary statistics from GWAS on INT and AD, International Age-related Macular Degeneration Consortium (IAMDGC) for sharing the summary statistics from GWAS on AMD, and the International Parkinson’s Disease Genomics Consortium (IPDGC) for sharing the summary statistics from GWAS on PD.

## References

1. Coleman HR, Chan CC, Ferris FL, 3rd, Chew EY. Age-related macular degeneration. Lancet. 2008 Nov 22;372(9652):1835–45.

2. Lim LS, Mitchell P, Seddon JM, Holz FG, Wong TY. Age-related macular degeneration. Lancet. 2012 May 5;379(9827):1728–38.

3. Mitchell P, Liew G, Gopinath B, Wong TY. Age-related macular degeneration. Lancet. 2018 Sep 29;392(10153):1147–59.

4. Turbert D. Top 5 risk factors for AMD. American Academy of Opthalmology; 2019; Available from: https://www.aao.org/eye-health/news/top-5-risk-factors-amd.

5. Woo SJ, Park KH, Ahn J, et al. Cognitive impairment in age-related macular degeneration and geographic atrophy. Ophthalmology. 2012 Oct;119(10):2094–101.

6. Xu W, Yu JT, Tan MS, Tan L. Cognitive reserve and Alzheimer’s disease. Mol Neurobiol. 2015 Feb;51(1):187–208.

7. Blume J, Rothenfusser E, Schlaier J, Bogdahn U, Lange M. Educational attainment and motor burden in advanced Parkinson’s disease - The emerging role of education in motor reserve. J Neurol Sci. 2017 Oct 15;381:141–3.

8. Raghavan NS, Vardarajan B, Mayeux R. Genomic variation in educational attainment modifies Alzheimer disease risk. Neurol Genet. 2019 Apr;5(2):e310.

9. Nalls MA, Blauwendraat C, Vallerga CL, et al. Expanding Parkinson’s disease genetics: novel risk loci, genomic context, causal insights and heritable risk. bioRxiv. 2019 2019-01-01 00:00:00.

10. Grover S, Del Greco MF, Stein CM, Ziegler A. Mendelian Randomization. Methods Mol Biol. 2017;1666:581–628.

11. Lee JJ, Wedow R, Okbay A, et al. Gene discovery and polygenic prediction from a genome-wide association study of educational attainment in 1.1 million individuals. Nat Genet. 2018 Jul 23;50(8):1112–21.

12. Fritsche LG, Igl W, Bailey JN, et al. A large genome-wide association study of age-related macular degeneration highlights contributions of rare and common variants. Nat Genet. 2016 Feb;48(2):134–43.

13. Grover S, Fabiola Del GM, Kasten M, Klein C, Lill CM, König IR. Risky behaviors and Parkinson’s disease: A Mendelian randomization study in up to 1 million study participants. bioRxiv. 2018 2018-01-01 00:00:00.

14. Savage JE, Jansen PR, Stringer S, et al. Genome-wide association meta-analysis in 269,867 individuals identifies new genetic and functional links to intelligence. Nat Genet. 2018 Jul;50(7):912–9.

15. Klein R, Klein BE, Jensen SC, Moss SE. The relation of socioeconomic factors to the incidence of early age-related maculopathy: the Beaver Dam eye study. Am J Ophthalmol. 2001 Jul;132(1):128–31.

16. Cackett P, Tay WT, Aung T, et al. Education, socio-economic status and age-related macular degeneration in Asians: the Singapore Malay Eye Study. Br J Ophthalmol. 2008 Oct;92(10):1312–5.

17. Zhang K, Zhong Q, Chen S, et al. An epidemiological investigation of age-related macular degeneration in aged population in China: the Hainan study. Int Ophthalmol. 2018 Aug;38(4):1659–67.

18. Grover S, Del Greco MF, Konig IR. Evaluating the current state of Mendelian randomization studies: a protocol for a systematic review on methodological and clinical aspects using neurodegenerative disorders as outcome. Syst Rev. 2018 Sep 24;7(1):145.

